# Automatic wound detection and size estimation using deep learning algorithms

**DOI:** 10.1101/2020.11.13.275917

**Authors:** Héctor Carrión, Mohammad Jafari, Michelle Dawn Bagood, Hsin-ya Yang, Roslyn Rivkah Isseroff, Marcella Gomez

**Affiliations:** Department of Computer Science, University of Puerto Rico, Río Piedras, SJ, PR; Department of Applied Mathematics, University of California, Santa Cruz, CA, USA; Department of Dermatology, University of California, Davis, Sacramento, CA, USA

## Abstract

Evaluating and tracking wound size is a fundamental metric for the wound assessment process. Good location and size estimates can enable proper diagnosis and effective treatment. Traditionally, laboratory wound healing studies include a collection of images at uniform time intervals exhibiting the wounded area and the healing process in the test animal, often a mouse. These images are then manually observed to determine key metrics —such as wound size progress– relevant to the study. However, this task is a time-consuming and laborious process. In addition, defining the wound edge could be subjective and can vary from one individual to another even among experts. Furthermore, as our understanding of the healing process grows, so does our need to efficiently and accurately track these key factors for high throughput (e.g., over large-scale and long-term experiments). Thus, in this study, we develop a deep learning-based image analysis pipeline that aims to intake non-uniform wound images and extract relevant information such as the location of interest, wound only image crops, and wound periphery size over-time metrics. In particular, our work focuses on images of wounded laboratory mice that are used widely for translationally relevant wound studies and leverages a commonly used ring-shaped splint present in most images to predict wound size. We apply the method to a dataset that was never meant to be quantified and, thus, presents many visual challenges. Additionally, the data set was not meant for training deep learning models and so is relatively small in size with only 256 images. We compare results to that of expert measurements and demonstrate preservation of information relevant to predicting wound closure despite variability from machine-to-expert and even expert-to-expert. The proposed system resulted in high fidelity results on unseen data with minimal human intervention. Furthermore, the pipeline estimates acceptable wound sizes when less than 50% of the images are missing reference objects.

**Author summary:** Knowledge of the wound size changes over-time allows us to observe important insights such as rate of closure, time to closure, and expansion events, which are key indicators for predicting healing status. To better perform wound measurements it is essential to utilize a technique that returns accurate and consistent results every time. Over the last years, collecting wound images is becoming easier and more popular as digital cameras and smartphones are more accessible. Commonly, scientists/clinicians trace the wound in these images manually to observe changes in the wound, which is normally a slow and labor-intensive process and also requires a trained eye. The clinical goal is to more efficiently and effectively treat wounds by employing easy to use and precise wound measurement techniques. Therefore, the objective should be devising automatic and precise wound measurement tools to be used for wound assessment. To this end, we leveraged a combination of various state-of-the-art computer vision and machine learning-based methods for developing a versatile and automatic wound assessment tool. We applied this tool to analyze the images of wound inflicted lab mice and showed that our developed tool automated the overall wound measurement process, therefore, resulting in high fidelity results without significant human intervention. Furthermore, we compared results to two expert measurements. We found variability in measurement even across experts further validating the need for a consistent approach. However, qualitative behavior, which is most important for predicting wound closure, is preserved.

## Introduction

A fundamental metric for studying wound healing is wound size and, generally it is the first criterion to be considered in a wound assessment process [1,2]. Several studies provide many reasons regarding the importance of wound measurement, such as monitoring the healing rate and progress, evaluating the effectiveness of treatments, and identifying the static wounds, to name a few [1–3]. In addition, plotting size changes over-time allows us to observe the rate of closure, time to closure, expansion events, and other insights which are key indicators for predicting healing status [4].

Different approaches are available to perform wound measurements and to be able to compare study outcomes, it is essential to employ a similar technique every time [2]. There exist many commercially available wound measuring cameras that utilize built-in algorithms for wound size measurements [5–7], however, not all the clinics or scientific labs have access to them due to high expenses. Collecting wound images is becoming easier and more popular as digital cameras and smartphones are more accessible and, therefore, increasingly being used as a means of recording/monitoring the wound healing progress. Commonly, scientists/clinicians reference these images manually to observe changes in the wound, which is normally a slow and labor-intensive process. This approach also requires a trained eye. However, this can quickly become a bottleneck as the experiment size grows and interesting big-data insights could be lost. Furthermore, these images are not always taken with the expectation of computerized image processing later down the road, so not much care is taken to capture clean, clear, and uniform data. While the images can provide greater information (e.g., tissue type, skin condition, etc.) about the wound, ensuring the consistency of the imaging is essential [1, 8].

This generally requires the following conditions among others, (i) to always have a ruler in the wound images, (ii) the distance between the wound and the camera should ideally be identical each time, and (iii) the angle of the wound relative to the camera lens should remain constant [8]. Considering the fact that satisfying all these requirements is not always possible, it is important to create a process that is robust against these variations. The potential tools for tackling these challenges are computer vision and machine learning-based methods. Among the state-of-the-art machine learning-based techniques, the deep learning algorithms are known for their performance against varied data distributions and speed if designed accurately. Therefore, leveraging the deep learning-based algorithms can enhance this task with higher accuracy sufficient to be used in the wound assessment process. The deep learning methods are expanding their popularity in the biological and medical applications and more specifically in image analysis, including but not limited to classification and segmentation [9–13].

Among the existing image-based techniques in wound assessment, most of them are focused on the classification of different wound types or conditions [14–18]. However, segmentation techniques, and more specifically semantic segmentation methods, are essential for developing a wound size measurement algorithm. There are several image-based wound periphery measurement methods which utilize deep and non-deep learning-based wound image segmentation approaches [20–27]. Deep learning-based techniques have the potential to be significantly more accurate but require larger datasets [19]. Additionally, many of the methods above aiming to accurately quantify wound features require customized data sets that include reference objects for size and color assessment or various imaging conditions to account for distortion. Finally, robustness to image quality or missing reference objects is rarely discussed. Improving on such methods will allow researchers to extract information from data sets not generated with AI algorithms in mind, thereby increasing usable data sets for analysis and understanding.

Here, we combine variations of two deep-learning techniques previously used for wound detection and segmentation. Our pipeline detects the wounded area using a deep learning-based object detection algorithm known as YOLO [28], where it allows us to automatically crop the wound-of-interest in unseen images at different resolutions, and leverages U-net as its main segmentation technique. However, we present a method of complimenting traditional techniques to develop a pipeline to automate wound size estimation robust to poor image quality and missing or deteriorated reference objects. We achieve this through strategic data augmentation and a custom post-processing algorithm. Our developed pipeline has the following advantages:

- With only 256 images in the data set, it produces high fidelity results without significant human intervention.
- It is robust against camera changes (e.g., distance, translation, angle).
- It is robust to wound occlusions and uneven lighting.
- Our algorithm can adapt to use any known size objects as a reference, in this work we used a ring-shaped splint around the wound as its reference object with known size.
- Applying several post-processing techniques, our pipeline can provide acceptable size estimate when approximately less than 50% of images are missing a reference object.
- Our customized object detection algorithm can accurately determine the wound-of-interest when multiple wounds are visible. This is done by choosing the detected wound with the highest confidence score.

Our pipeline generalizes the overall process, therefore, resulting in a more precise estimation. This method is user friendly in that it can be used by a scientist with minimal consideration during data generation. Finally, we compare our results to that of experts and show preservation of information relevant to predicting wound closure and other clinical studies. We note that discrepancies in wound segmentation can occur across experts. This further motivates the need for a consistent method to quantify wound size but also sheds light on the challenge in getting consistency in annotated data. For consistency, all manually segmented wounds in the generation of training data were done by a single person.

## Materials and methods

Automatic wound size calculations were performed using a custom-designed deep learning-based pipeline (see Fig. 1) that intakes mice wound images (splinted or missing/damaged splint) and extracts relevant information from wound images such as wound-of-interest location and wound periphery size estimates. We developed customized algorithms based on the well-known object detection and medical instance segmentation deep learning algorithms known as YOLO [28] and U-Net [10], respectively, and adapted them to our particular task. A preprocessing task was performed to make images more consistent and easier to work with. Detection and location of the wounded area were performed by a YOLOv3 [29] based object detection algorithm trained on our dataset, where we manually labeled bounding boxes around the wound for each image on our training set. This architecture was chosen because the output annotations are scale-invariant and will allow us to crop images of any resolution. Cropping was done by an algorithm able to read-in output annotations and crop around the detected midpoint of the wound-of-interest. Size estimation was performed by developing an image segmentation algorithm aimed at segmenting the wounded and ring area (i.e., the inner area of the splint located around the wound) for each wound image. Some post-processing was performed to clean up the segmentation results ensuring a better estimation of the wound size. Further explanations are provided in the following subsections.

**Fig 1.**
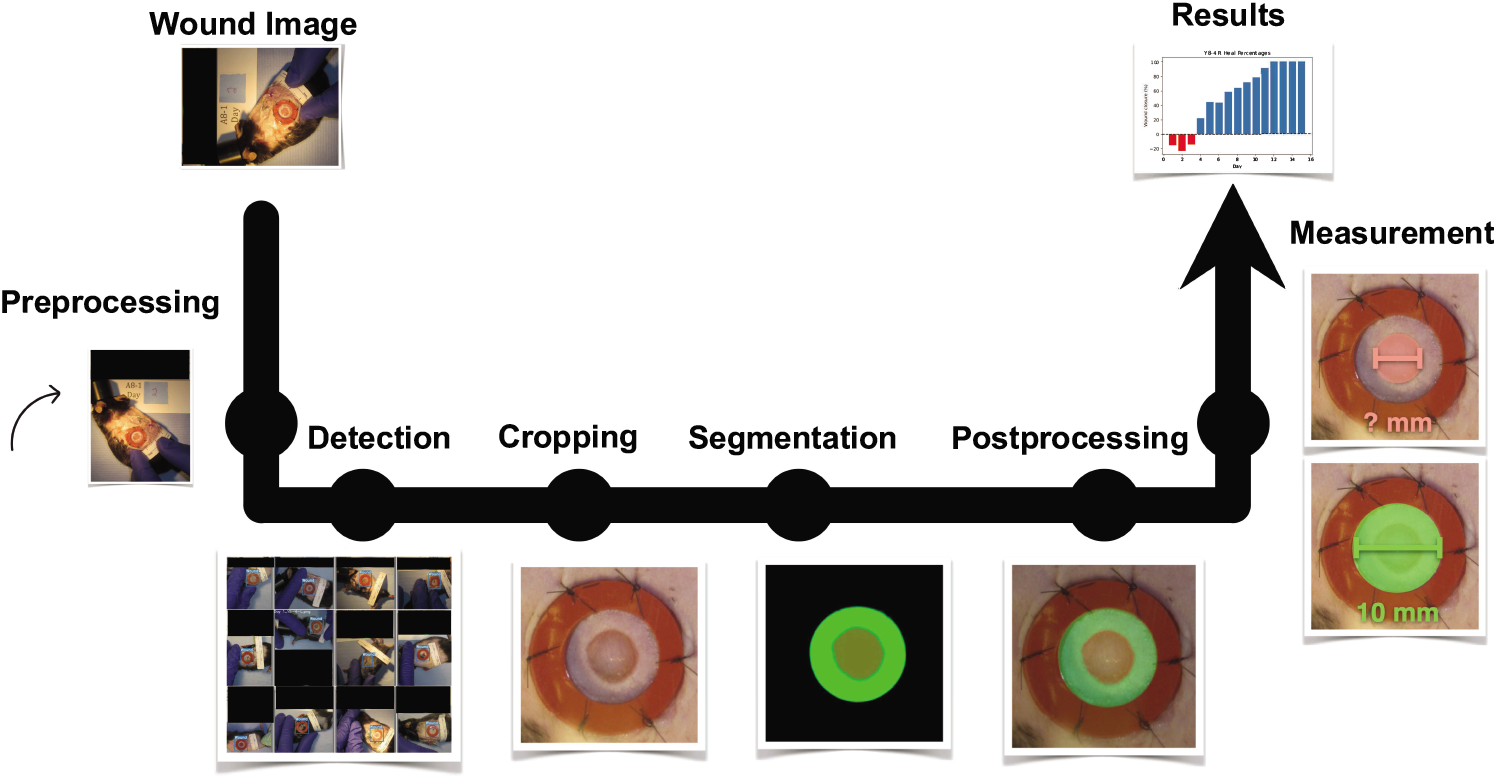
Custom-designed end-to-end deep learning-based image analysis pipeline. The pipeline consists of 6 major blocks for performing the preprocessing, detection, cropping, segmentation, postprocessing, and measurement. <u>Note: All the black boxes have been added later to obscure the protocol number and any animal identifying information.</u>

### Wounding surgery and daily imaging

The mouse wounding surgery and the animal protocol was approved by UCD IACUC. Two cohorts of C57BL/6J mice (male, Jackson lab, Sacramento, CA) were used in 2 independent experiments. One day prior to wounding, dorsal surfaces of mice were shaved and then depilated (Nair, Church & Dwight Co., Inc., Ewing, NJ). On day 0, mice were anesthetized with 2.5% isoflurane and injected subcutaneously with analgesic (Buprenex, 0.05 *mg/mL*, 2 *μL/g* body weight). After betadine and alcohol washes of the mouse dorsum, two sterile silicon splints (10 *mm* inner diameter, 16 *mm* outer diameter, 1.6 *mm* thick) were glued using cyanoacrylate (Scotch® Super Glue, 3*M* ™, Maplewood, MN) to the back skin 35 *mm* from the base of the skull and 10 *mm* on either side of the spine. Splints were further secured with six interrupted sutures (Ethilon 6 − 0 nylon suture, #697*G*, Ethicon, Bridgewater, NJ). Wounds were created in telogen skin, evidenced by the absence of visible dark follicles. All mice received a 6 − *mm* full-thickness, circular wound in the center of each splint (2 total wounds per mouse) by a biopsy punch (Miltex® Instruments, Integra Lifesciences, Princeton, NJ) and surgical microscissors. A 16 *mm* circular plastic coverslip was applied on top of the splints and the entire dorsum area was sealed with a transparent, semi-permeable dressing (Tegaderm™, 3*M* ™, Maplewood, MN) adhered with liquid bandage (New Skin®, Advantice Health, Cedar Knolls, NJ). The mice were housed singly after wounding, and daily mice were weighed and wounds were imaged. On day 15 post wounding, animals were euthanized by exsanguination and tissues were collected for analysis detailed below. The wounds were photographed using the digital camera application on the iPhone X that was attached to a tripod using a clamp to ensure the camera was at the same height for every photograph taken over the 16 days.

### Dataset

The original dataset consists of approximately 256 images of wound inflicted lab mice (see “Wounding surgery and daily imaging” for details). It covers four mice in Cohort 1 (C1) and four mice in Cohort 2 (C2) over 16 days of healing (i.e., starting from day 0 (surgery) to day 15). The two cohorts represent two distinct experimental conditions and so the results are grouped by cohort. Each mouse has two wounds, one on the left and the other one on the right side. Each wound has a ring-shaped splint surrounding the wounded area, where its inner diameter is ≈ 10*mm* and its outer diameter is ≈ 16*mm*. Since the splint has a fixed inner/outer diameter, it is used as a reference object with known size in our pipeline. It should be noted that these splints are missing or have substantial damage for over 25% of the images in this dataset. There are several challenges present in these images, for example, there are variations in image rotation (landscape vs portrait), lighting conditions (Fig. 2A, 2B), unreliable or occluded tape measure placement (Fig. 2C, 2D), significant occlusion of the wound (Fig. 2E, 2F), multiple visible wounds (Fig. 2G, 2H) and wound location relative to the image frame. These challenges pushed the research towards a combination of deep learning algorithms and traditional image processing methods. We split the dataset into a train (75%), validation (12.5%), and test (12.5%) subsets while making sure the aforementioned challenges are well represented across each cohort. Additionally, we made sure there were images from each mouse and both their sides on each split.

**Fig 2.**
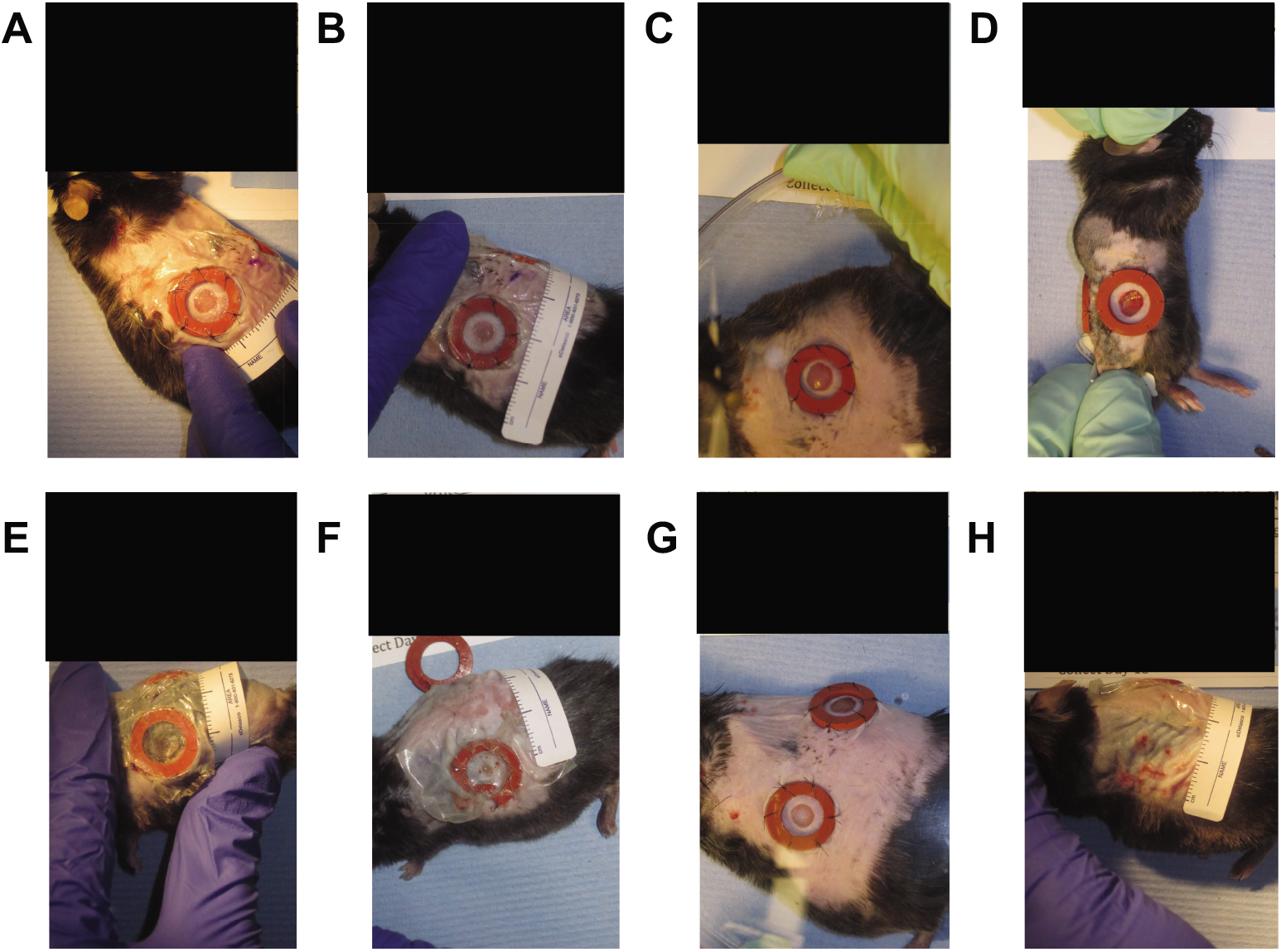
Dataset Challenges: Images were sometimes directly illuminated by a warm tone light source (**A**) and other times they were not (**B**). The measuring tape is often placed outside the frame or occluded (**C, D**). The wound surface area can often become significantly occluded by plastic coverings, reflections, debris, etc (**E, F**). There are two wounds on each mice, it is possible for an image intended to observe only the left wound contains a clear instance of the right wound (**G**). It is also possible that stitches holding the splint in place inflict secondary wounds around the wound of interest (**H**). <u>Note: All the black boxes have been added later to obscure the protocol number and any animal identifying information.</u>

### Manual Measurement

#### Manual Measurement (Periphery)

For the purposes of training our segmentation algorithm we defined wound periphery as the boundary furthest from the wound center at which the skin appearance changes from normal to abnormal (e.g., the boundary between “Region **A**” and “Region **B**” in Fig. 3). This kind of wound/skin boundary segmentation is common among computational wound segmentation works [19–23]. Given our automatically generated cropped wound dataset (see “Detection” Section for details) we used the open-source software Labelme [30] to draw polygons around the wound edge for all the images in our dataset. We also drew polygons around the splints (when available) as these manual measurements also serve as ground-truth annotations for our Training and Validation sets (Fig. 4). These measurements may not be pixel-perfect, especially for the more challenging images in the dataset where the wound edge is ambiguous (see Fig. 6). To ensure uniformity of the annotation, the same non-expert performed all the manual annotation of raw data from scratch using a consistent protocol.

**Fig 3.**
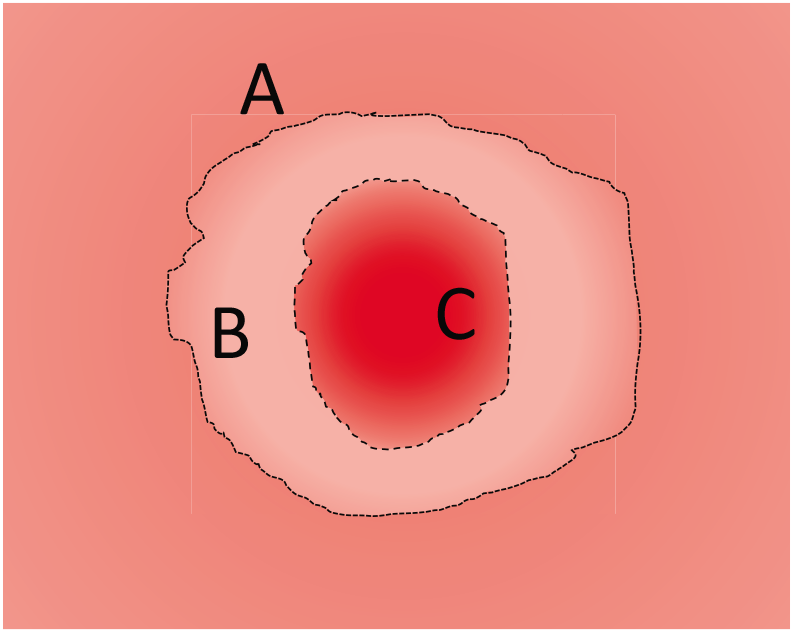
Cartoon image showing different wound area: “Region **A**” shows normal skin. “Region **B**” shows the regeneration of epidermis. “Region **C**” shows wound center.

**Fig 4.**
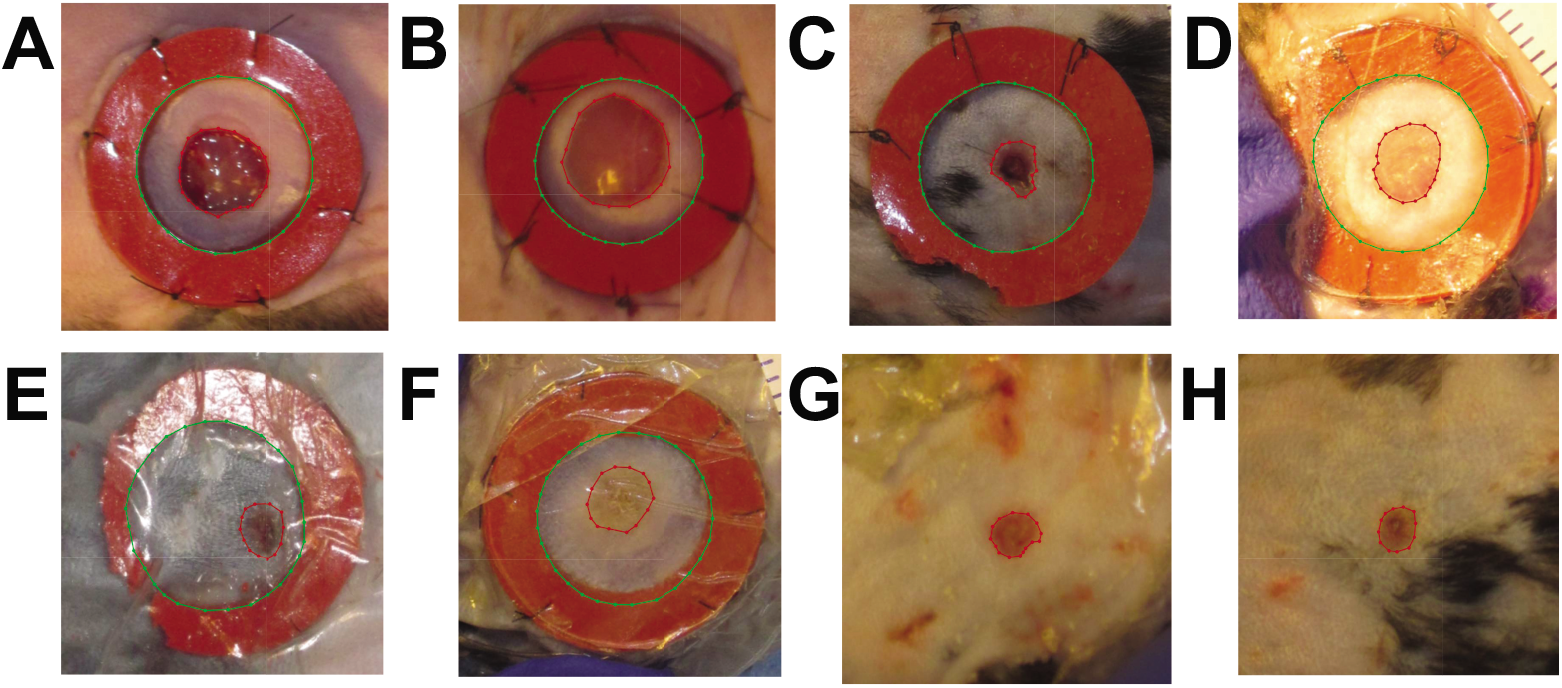
Wound Periphery Annotations: Red lines delineate the wound while green lines delineate the splint. Early day zero images (**A, B**). Midway images (days 4-8) with splint (**C, D, E F**) and without splint (**G, H**).

#### Manual Measurement (Expert)

For comparison, we describe here the method used by selected experts to delineate the wound edge. Wound areas were macroscopically measured on daily images (see “Wounding surgery and daily imaging” and “Dataset” for details) using ImageJ software, version 1.8.0_172 (http://imagej.nih.gov/ij). Briefly, the freehand selection tool was used to trace the neo-epithelial/granulation tissue border (e.g., the boundary between “Region **B**” and “Region **C**” in Fig. 3). Identifying the neo-epithelial/granulation tissue border from wound images can be challenging and delineations of said borders may vary across experts. More accurate analysis could be done by utilizing other types of image analysis (e.g., fluorescence excitation images and histology images), techniques that have inherent disadvantages such as the need for in vivo imaging (for fluorescently labeled markers) or terminal experiments (histological images) precluding temporal analysis [31, 32]. The pixel numbers of the wound areas was obtained using the measure function in ImageJ. Then the percentage of wound area closed relative to the original wound was calculated using the following formula: Relative closed wound area (%) = [1 – (open wound area/original wound area)] × 100. Wound closure was complete (100% closed) when the entire wound surface area was covered with neo-epithelial tissue.

### Pre-processing

To make the input images more consistent and easier to work with, the original images were automatically rotated to portrait orientation (i.e., vertically rotated). This task is accomplished by reading the width and height of the input images and performing a 90-degree rotation if the width is larger than the height of the image. The images are also down-sampled to half of their original size, which facilitates handling of the data, the training process and does not sacrifice image quality to a significant degree.

### Detection

To automatically generate the crops, we manually labeled bounding boxes around the wound (including the splint when available) for each image on our training set using the open-source software Labelme [30] and then trained a YOLOv3 based object detection algorithm (see Fig. 5A). The YOLO output includes scale-invariant coordinates denoting the detected object midpoint, thus, given these coordinates, a custom-designed algorithm crops the original high-resolution images to produce 352 × 352 wound-of-interest sub-images. In addition, we customized the YOLO output file to include a confidence score for the cropping algorithm to choose the detected object with higher confidence score as the wound-of-interest when multiple wounds are detected (see Fig. 5B). This experiment generated our consistent cropped wound dataset used from this point forward. This new dataset surfaced previously hard-to-notice challenges such as different camera angles (Fig. 6A, 6B), camera focusing issues (Fig. 6C, 6D), hair growth (Fig. 6E, 6F) and day zero size differences (Fig. 6G, 6H).

**Fig 5.**
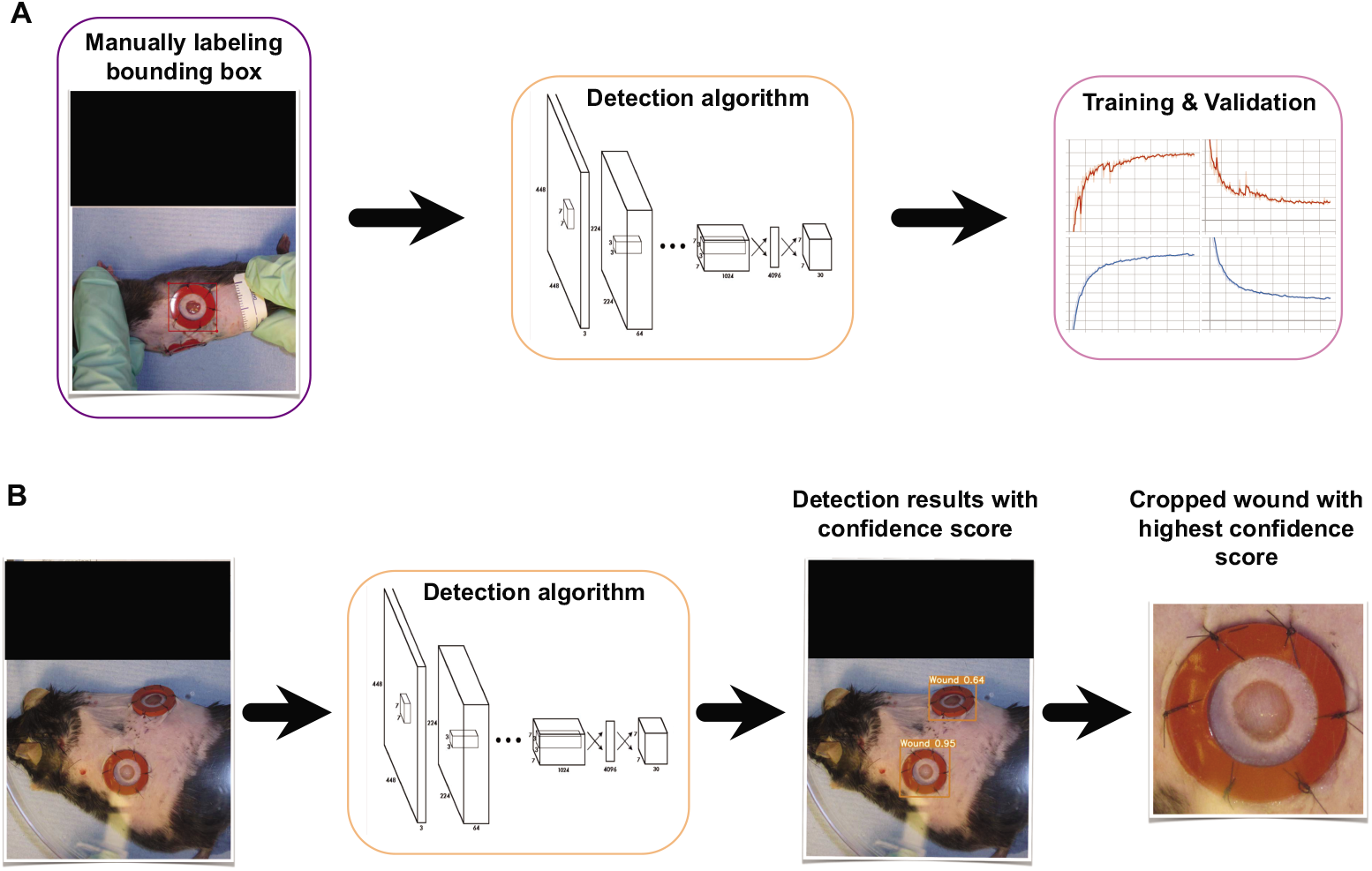
Details of detection and cropping process. A) Manual labeling bounding box, Training, and Validation, B) Testing (i.e., detecting the wound-of-interest and cropping the wound with highest confidence score. <u>Note: All the black boxes have been added later to obscure the protocol number and any animal identifying information.</u>

**Fig 6.**
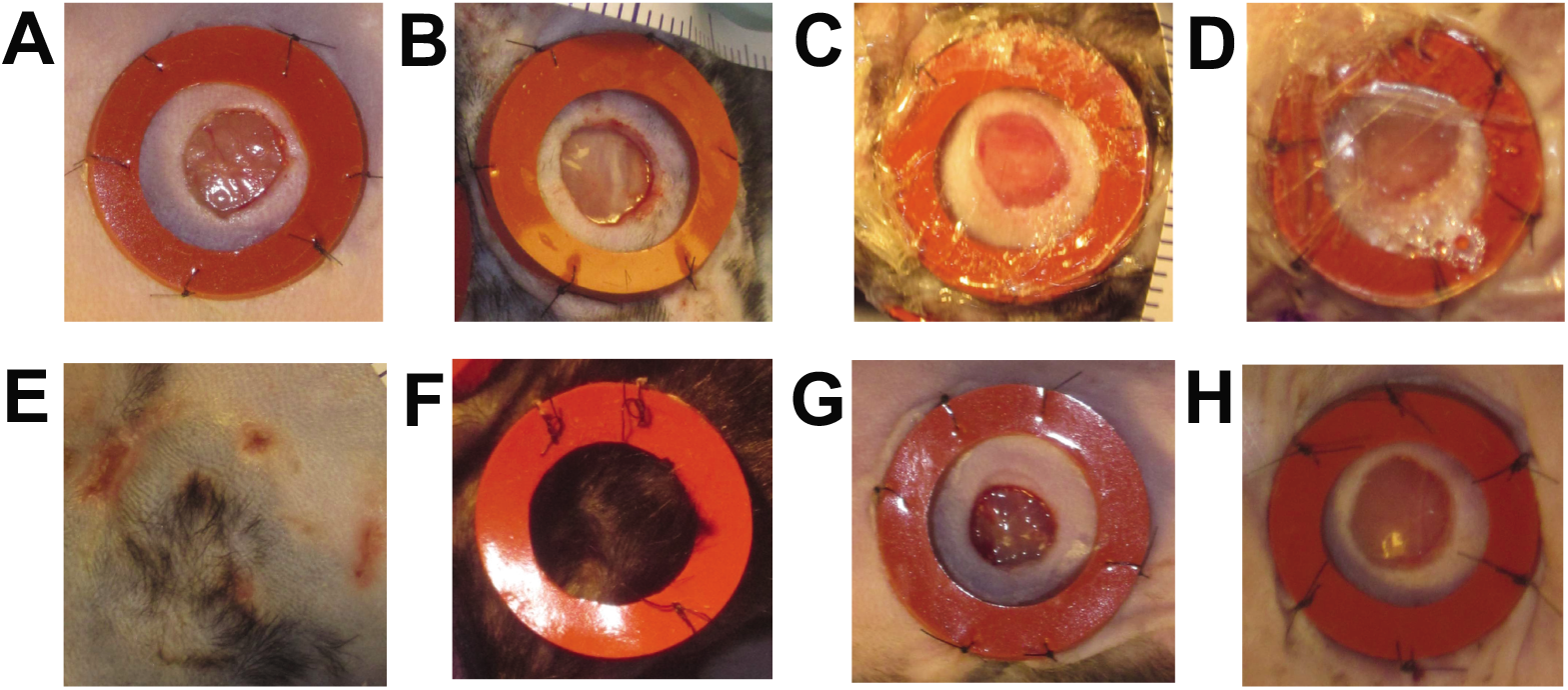
New Dataset Challenges: Camera can appear to be at unknown angles relative to the surface of the wound, 90° (**A**) unknown angle (**B**). Camera is often not focused at the wounded area or the Tegaderm dressings are still there, introducing significant blurriness (**C, D**). Hair may grow back before the end of the experiment (i.e., before the complete wound closure), occluding the wounded area **(E, F)**. Initially, we believed the day zero wound sizes to be the same across all mice given that the same tool was used to produce the wounds but close inspection seems to show variability **(G, H)**. <u>Note: the images shown here are a direct output of our automatic cropper, no alterations have been made.</u>

### Segmentation

To observe the different healing rates and generate wound periphery size estimates where wound periphery is defined as the boundary at which the skin appearance changes form normal to abnormal, we developed an image segmentation algorithm. The task for this model is to segment the wounded area and ring area (i.e., the inner area of the splint located around the wound) for each wound image. Given these masks, we could then estimate the unknown wound area from the known ring area (see “Wound Area Calculation” section). To perform this, we trained a U-Net [33] based algorithm on our cropped dataset, where we manually annotated our training set by drawing polygons around both wound areas and the inner ring (whenever the splint was available) (see Fig. 7A). To boost our available training data, we applied heavy data augmentation such as vertical and horizontal flips (performed at random 50% of the time), rotation (applied randomly from a −90 to 90 degree range. This means we apply affine rotation on the y-axis to input data images by a random value between −90 and 90 degrees), linear contrast (applied at random from a 0.50 to 1.50 range. This means we modify the contrast of images according to 127 + alpha*(v-127), where v is a pixel value and alpha is sampled uniformly from the interval [0.5, 1.5] (once per image)), pixel intensity (multiplier applied at random from a 0.50 to 1.50 range. This means we multiply all pixels in an image with a specific value [0.5, 1.5], thereby making the image darker or brighter), and Gaussian blur (applied 35% of the time at a random severity sigma inside a 0 to 6 range). We are using the “On The Fly Data Augmentation” and the generated instances change at each epoch. There are 6 different augmentation techniques that are applied at random for each training sample. We trained for 1000 epochs at a batch size of 4, therefore the size of the training data after augmentation could be up to ≈ 4000. It is worth noting that data augmentation is only applied in the training phase. After successfully training and validation of the segmentation algorithm, we perform testing on the unseen images (Fig. 7B). Figure 7 illustrates the overall segmentation process (i.e., manual annotation, training, validation, and testing) in detail. Given the masks obtained from the segmentation algorithm (see Fig. 7B), we estimated the unknown wound area from the known ring area (details are provided in the Post-processing and Automatic Measurement sections).

**Fig 7.**
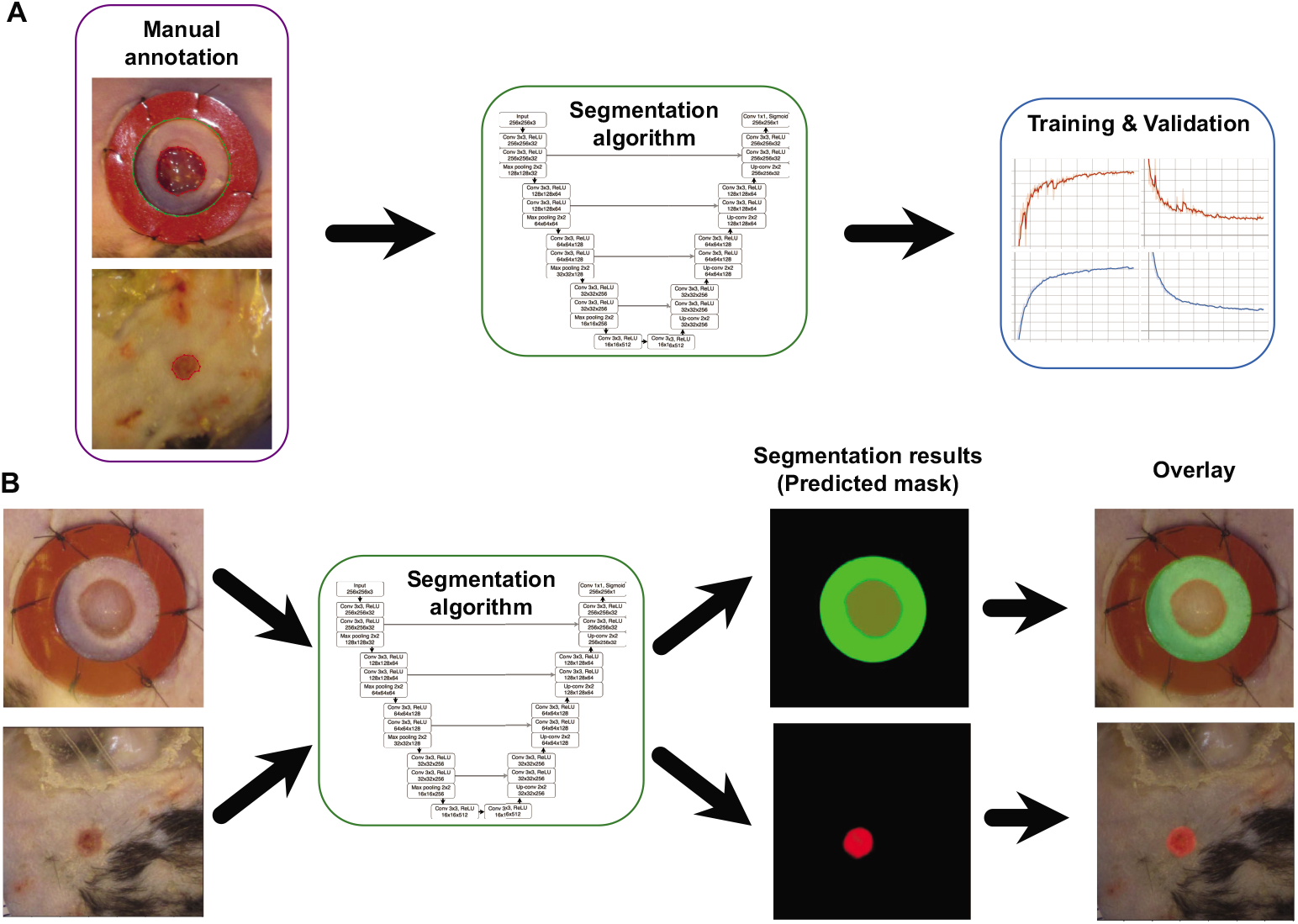
Details of image segmentation process. A) Manual annotation, Training, and Validation, B) Testing (i.e., mask prediction).

The results achieve 0.96 Dice and 0.93 IoU scores (see Fig. 8) and generating good predictions on the test set. The Dice score tends to measure closer to the average performance while the IoU score tends to measure closer to worst-case performance. Dice is more commonly used for segmentation. The blue plots in Fig. 8 show the training loss and training accuracy (i.e., dice and IoU coefficients) and the red plots show the validation loss and validation accuracy (i.e., dice and IoU coefficients). The loss graph shows that the loss is gradually decreasing toward zero and then stabilized. The dice and IoU coefficients show that the accuracy is gradually increasing toward 100% and then remain steady. This means the performance of the model is promising. Some post-processing was later applied to clean up the predictions.

**Fig 8.**
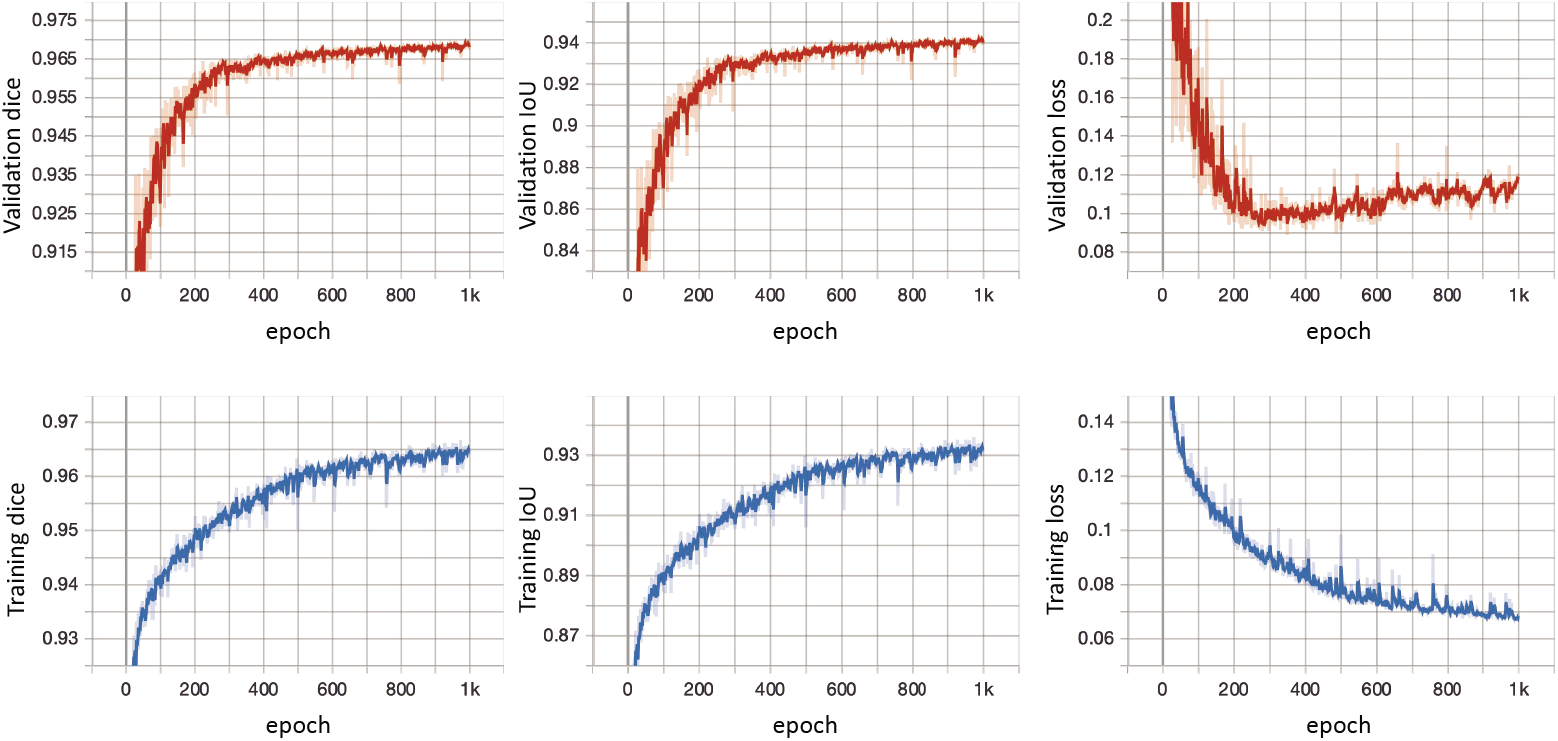
Training and Validation curves, including loss, dice, and IoU. The blue plots show the training loss and training accuracy (i.e., dice and IoU coefficients) and the red plots show the validation loss and validation accuracy (i.e., dice and IoU coefficients). The loss graph shows that the loss is gradually decreasing toward zero and then stabilized. The dice and IoU coefficients show that the accuracy is gradually increasing toward 100% and then remain steady.

### Post-processing

The post-processing involved traditional image processing techniques. To clean up the predicted ring mask, we select the largest connected component. To account for false-positive wounds masks (i.e., wounds that can be present in an image but are not of interest) we select the connected component that is closer to the middle of the cropped image within a certain threshold in position from the center. In other words, we selected the connected component that is closest to the center-point of the cropped image (i.e., the pixel at (*x* = 176, *y* = 176)). This technique eliminates secondary splint wounds developed around the outer perimeter of our wound of interest. To estimate size when no ring is detected, we overlay the average ring mask calculated from the whole set or we search for the closest known size wound with a ring detection and apply that ring mask to calculate the size of the unknown wound. To clean any missing points in our detections we insert the average between the previous and next points. These techniques are summarized as follows:

1. To clean up the ring mask, we select the largest connected component.
2. To clean up the wound mask we select the connected component that is closer to the middle (over a certain size).
3. To estimate size when no ring is detected, we overlay the average ring mask calculated from the whole set.
4. Alternatively, to estimate wound size when no splint is detected, we perform a search for the closest sized wound with a detected ring and apply that ring mask to estimate size.
5. To clean any missing points in our detections we insert the average between the previous and next points.

It is worth noting that, when the dataset contains many unsplinted wounds, it is beneficial to trade the inference speed of technique 3 for the more accurate estimates of technique 4.

### Automatic Measurements

Having the wound images detected and segmented properly, various analysis is performed to extract meaningful information. Here, we calculated the wound area and wound closure percentage.

#### Wound Area Calculation

Wound area calculation was performed using the following equation:

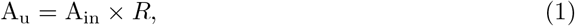

where A_u_ is the unknown wounded area of interest, A_in_ is the known inner ring area, and *R* is the ratio of wounded area to inner ring pixels.

#### Wound Closure Calculation

Wound closure percentage calculation was performed using the following equation:

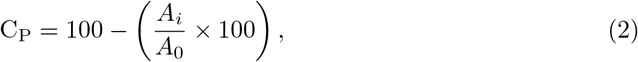

where C_P_ is the wound closure percentage, *A*_0_ is the wounded area at day 0, and *A_i_* is the area of the wound at day *i*. Day 0 size is considered 0% closure, an increase/decrease in the size is considered as negative/positive percentage change, and the 100% closure means the wound is completely closed.

## Results

This section presents the extracted results by employing the developed pipeline to the experimental data of 8 different mice (see “Dataset” section for details). It is possible to perform different analyses using the extracted information. Here, we plotted the estimated wound closure percentage of each wound image over time. In addition, to have a better insight into the performance of the developed pipeline, we plotted the wound closure percentage using the measurements from our manually annotated wound and ring areas, noting that these manual measurements were performed by an individual unfamiliar with the experiment, and not a trained expert.

Figure 9 shows manual (periphery) and automatic measurements of average wound closure percentages for C1 mice (see Fig. 9 left-plot) and C2 mice (see Fig. 9 right-plot). The orange line shows the average wound closure percentage measurements based on the manual periphery annotations and the blue line shows the average wound closure percentages based on the automatic estimation using the developed pipeline.

**Fig 9.**
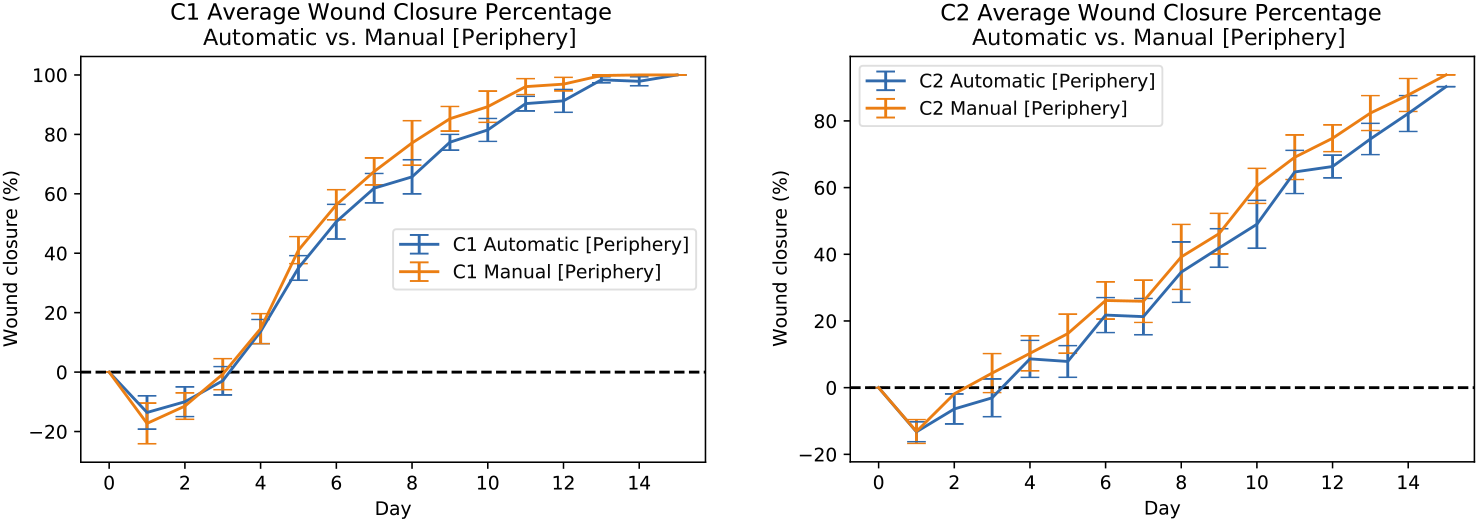
Automatic and manual measurements of wound periphery for C1 (left-plot) and C2 (right-plot) mice. The orange line shows the average wound closure percentage measurements based on the manual periphery annotations and the blue line shows the average wound closure percentages based on the automatic estimation using the developed pipeline.

Figures (S1 Fig and S2 Fig) show the wound closure percentages for C1 mice both left and right wounds. The results in Fig.(S1 Fig) are obtained based on the automatic estimation using the developed pipeline and the results in Fig.(S2 Fig) are obtained by using the measurements based on the manual periphery annotation of the images. In both figures, the left plot shows the wound closure percentages for the wound on the left side of 4 different C1 mice from day 0 to day 15 in different colors, and the right plot shows the wound closure percentages for the wound on the right side of those mice. Figures (S3 Fig and S4 Fig) show the same measurements but for C2 mice.

In order to further demonstrate the robustness of the developed pipeline and its various post-processing techniques dealing with a missing reference object, we artificially removed the splint masks for up to 50% of images. Figures 10 and 11 show the results of applying post-processing technique 3 and technique 4 respectively. These results show that the developed pipeline can provide an acceptable size estimate when approximately less than 50% of images are missing a reference object. In both figures (i.e., Figs. 10 and 11), the blue line shows the average wound closure percentages based on the automatic estimation using the developed pipeline for dataset having 75% (baseline) of the splints, the orange shows the same results but for having 60% of the splints, and finally green shows the same results but for 50% missing splints.

**Fig 10.**
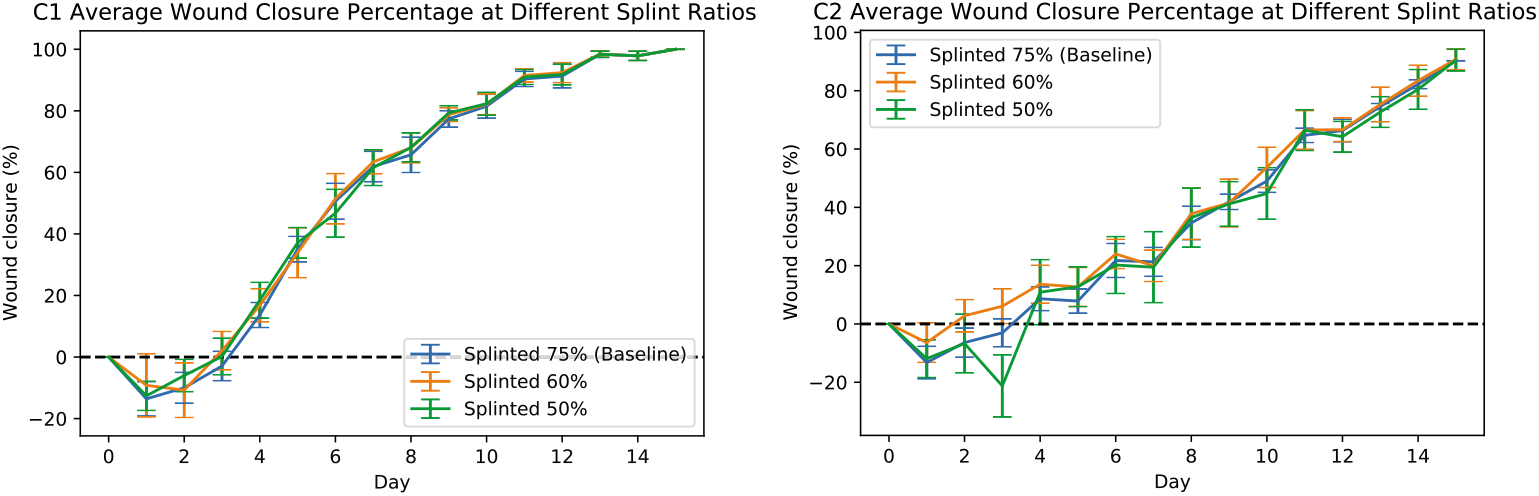
Automatic measurements of wound periphery using post-processing technique 3 for C1 (left-plot) and C2 (right-plot) mice when 75% (baseline), 60%, and 50% of the images have splints. The blue line shows the average wound closure percentages based on the automatic estimation using the developed pipeline for dataset having 75% (baseline) of the splints, the orange shows the same results but for having 60% of the splints, and finally green shows the same results but for 50% missing splints.

**Fig 11.**
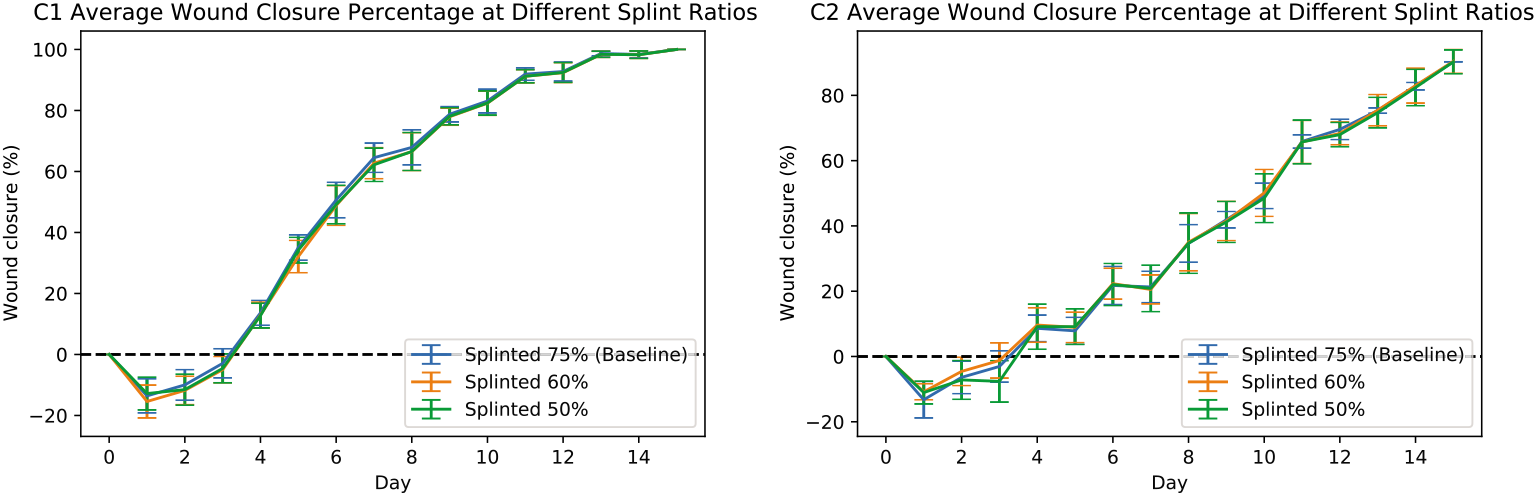
Automatic measurements of wound periphery using post-processing technique 4 for C1 (left-plot) and C2 (right-plot) mice when 75% (baseline), 60%, and 50% of the images have splints. The blue line shows the average wound closure percentages based on the automatic estimation using the developed pipeline for dataset having 75% (baseline) of the splints, the orange shows the same results but for having 60% of the splints, and finally green shows the same results but for 50% missing splints.

Furthermore, to demonstrate the robustness of the developed pipeline and its various post-processing techniques we consider a missing reference object at biased time points.

We artificially removed the splint masks of all the images in a fixed window of size 8 days (i.e., 0 – 7, 1 – 8, 2 – 9, 3 – 10, 4 – 11, 5 – 12, 6 – 13, 7 – 14, and 8 – 15). Figure 12 shows the results of applying post-processing technique 4 dealing with a biased missing reference object only for three different windows for better illustration. However, the measurements for all the different windows are available in (S1 File). In Fig. 12), the blue line shows the average wound closure percentages based on the automatic estimation using the developed pipeline for dataset having 75% (baseline) of the splints, the orange shows the same results but for those with all the splints removed for days 0 – 7, the green shows the same results but for those with all the splints removed for days 4 – 11, and finally red shows the same results but for those with all the splints removed for days 8 – 15. These results show that the developed pipeline can provide an acceptable size estimate.

**Fig 12.**
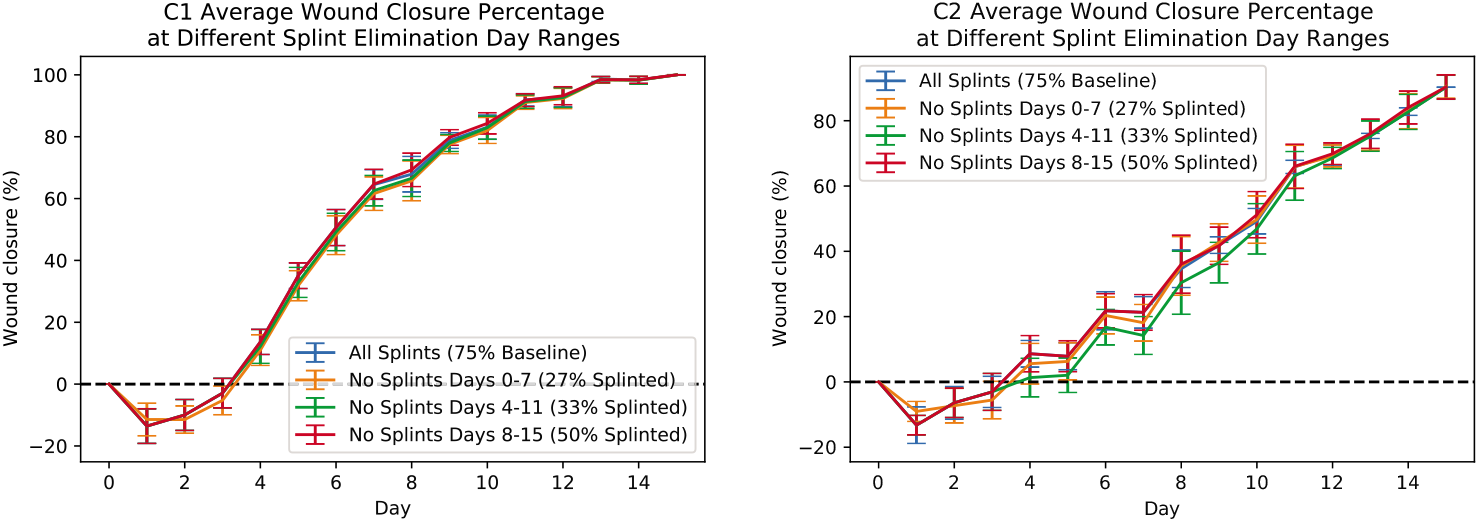
Automatic measurements of wound periphery using post-processing technique 4 for C1 (left-plot) and C2 (right-plot) mice dealing with a biased missing reference object. The blue line shows the average wound closure percentages based on the automatic estimation using the developed pipeline for dataset having 75% (baseline) of the splints, the orange shows the same results but for those with all the splints removed for days 0 − 7, the green shows the same results but for those with all the splints removed for days 4 − 11, and finally red shows the same results but for those with all the splints removed for days 8 − 15.

Moreover, to demonstrate the robustness of the developed pipeline and its various post-processing techniques to camera angle, we changed camera angle by 10 − 15 degrees for all the four mice in Cohort 1 (C1) and four mice in Cohort 2 (C2) in day 1, and statistically compared these measurements to measurements where camera angle was measured at 0 degree. Figure (S6 Fig) shows the Root Mean Square Error (RMSE) of estimated size of angled images comparing to those taken at 0 degree where the mean of the RMSE for all mice is 1.8088 and the median is 1.3472. To further study whether the RMSE of the estimated results for the angled images are acceptable, we calculated the RMSE of images (in the original dataset) taken at Day 1 comparing to those taken at Day 0. Figure (S6 Fig) shows the results where the mean of the RMSE for all mice is 4.1746 and the median is 4.275. Figure 13A shows the RMSE for all mice in the angled images dataset and those in the original dataset comparing Day 1 to Day 0. There exist one outlier in the RMSE of the angled images where Fig. 13B and 13C show the raw image and its masks. It is worth noting that it is hard for human annotators to analyze this image.

**Fig 13.**
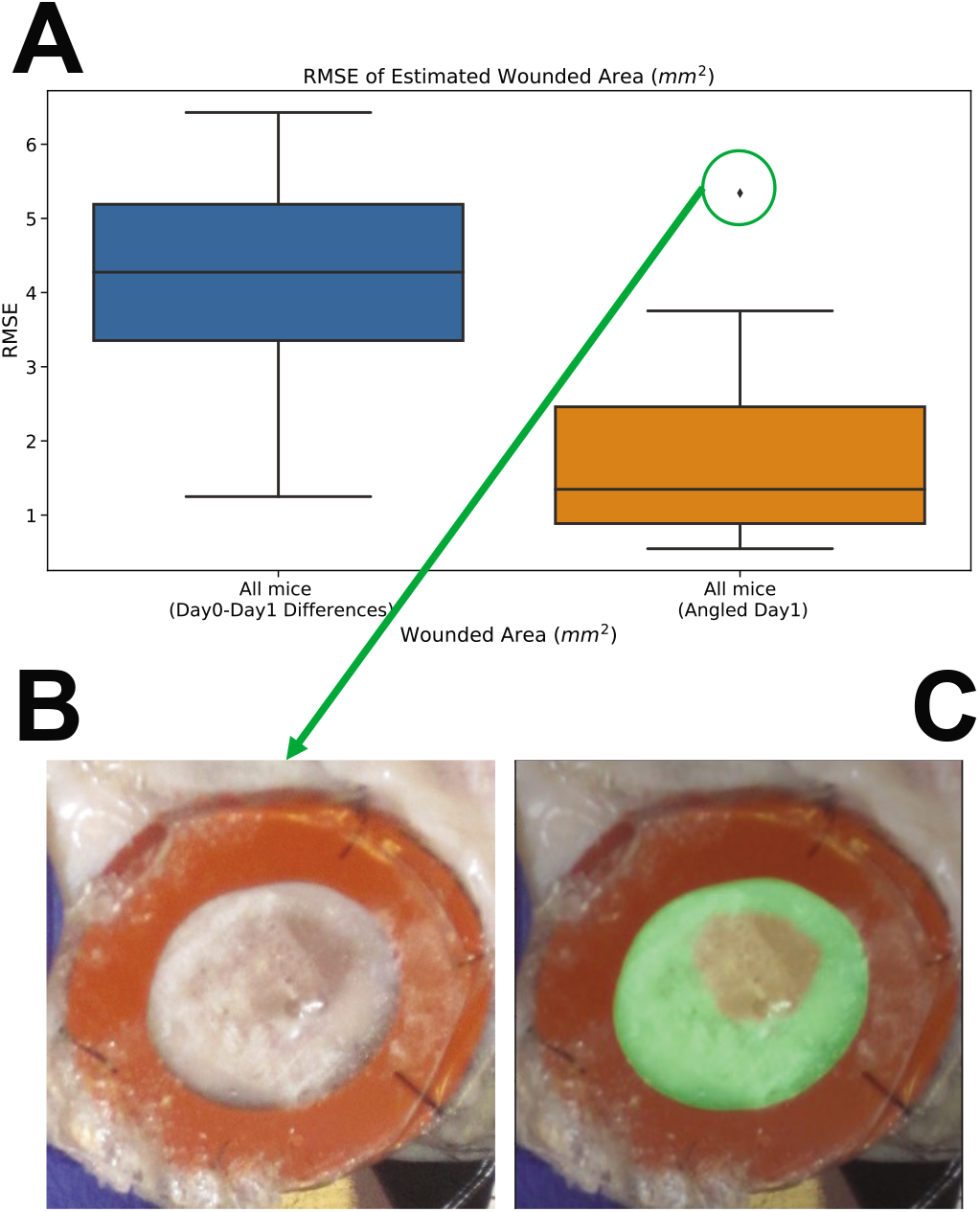
Comparison of the results for automatic analysis of wound periphery for wounds with changed camera angle by 10-15 degrees with those in the original dataset. (A) shows the RMSE for all mice in the angled images dataset and those in the original dataset comparing Day 1 to Day 0. (B,C) show the raw image and its masks for the outlier in the RMSE of the angled images.

Finally, to demonstrate the robustness and generalizability of the developed pipeline and its various post-processing techniques to dealing with wounds with different initial sizes, we employed the developed pipeline to analyze those wounds with the initial size of ⪆ 8 – *mm*. Figure 14 shows the results of applying the developed pipeline for analyzing two different images with different initial sizes. These results show that the developed pipeline can provide accurate masks for both wounded area and inner area (i.e., accurate wound estimation). It worth to note that, we used the same pipeline with no change in its parameter to perform these analysis.

**Fig 14.**
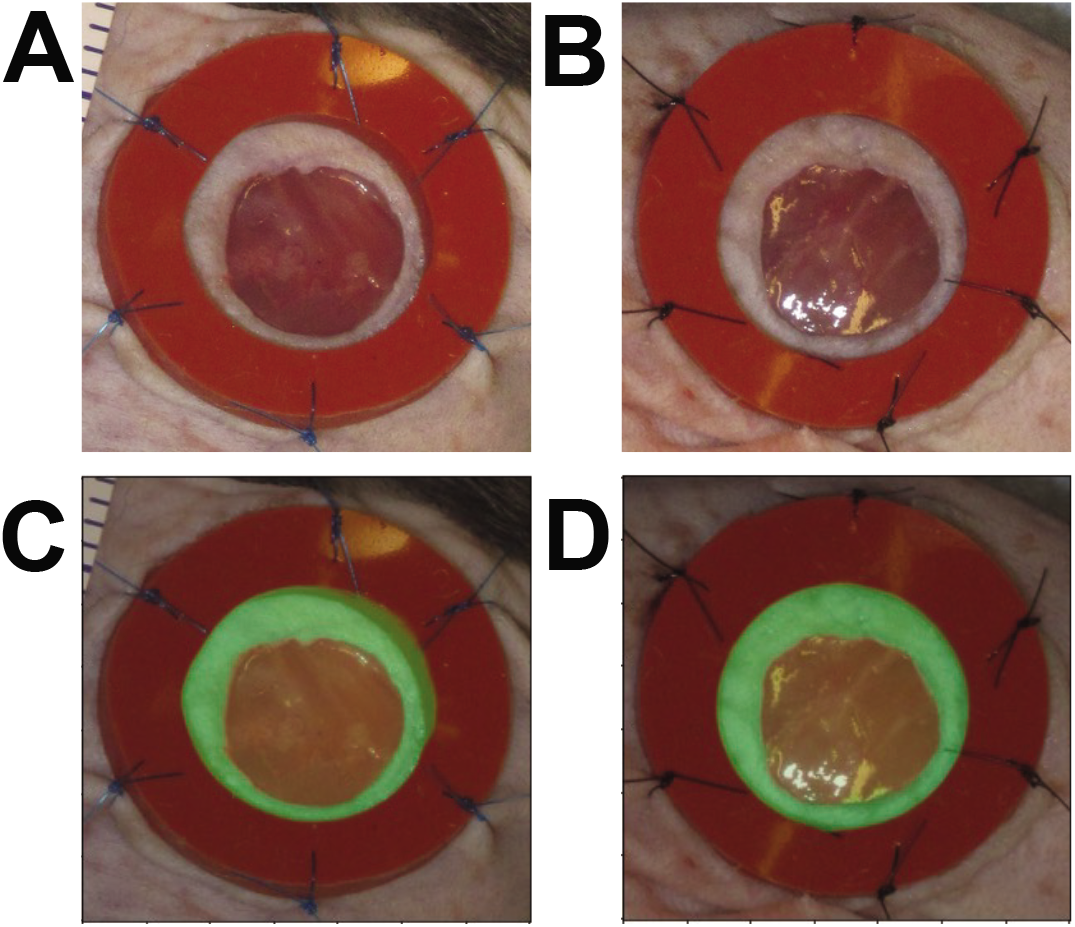
Automatic analysis of wound periphery for wounds with different initial sizes using developed pipeline. The raw image and its masks for sample 1 (A,C) and sample 2 (B,D). Results show that the developed pipeline can provide accurate masks for both wounded area and inner area.

## Discussion

Measuring the wound size is an indispensable part of the wound assessment process where its precision directly impacts the proper diagnosis and effective treatment. Here, we demonstrated that our developed pipeline could automatically measure laboratory wounds, therefore, can be beneficial for ultimately translating to the human wound measurement. We have shown that leveraging the deep learning-based algorithms can enhance this task with higher accuracy sufficient to be used in the wound assessment process. Our developed pipeline has several advantages over standard computer vision techniques. It provides high fidelity results with minimal human intervention. It is robust against camera changes (e.g., distance, translation, angle). It automates the whole measurement process which is a time-consuming and laborious task if manually performed. Thanks to the several post-processing techniques employed in our pipeline, it provides an acceptable size estimate when approximately less than 50% of images are missing reference object. Furthermore, Our pipeline detects and localizes the wounded area and it allows us to automatically crop the wound-of-interest in unseen images at different resolutions. In addition, our customized detection algorithm could accurately determine the wound-of-interest when multiple wounds are visible. It utilized the splint around the wound as its reference object with known size. In addition, our algorithm could use any known sized object in the image as the reference. Ultimately, generalizing the overall process, it results in a more precise estimation.

When performing the tracing, defining the wound margins could be subjective and sometimes difficult because the wound margins are not always clearly visible [2]. Therefore, the choice of wound margins can vary from one individual to another even among experts. Figure 15 shows the comparison of three different tracings (i.e., Manual (periphery) vs. Manual (experts (#1 and #2))). Fig. 15A, 15B, and 15C show similar trends in tracing performed by two different experts (#1 and #2) and a non-expert for the same wound (taken before day 3) respectively. Fig. 15D, 15E, and 15F show different trends in tracing performed by two different experts (#1 and #2) and a non-expert for the same wound (taken after day 3) respectively.

**Fig 15.**
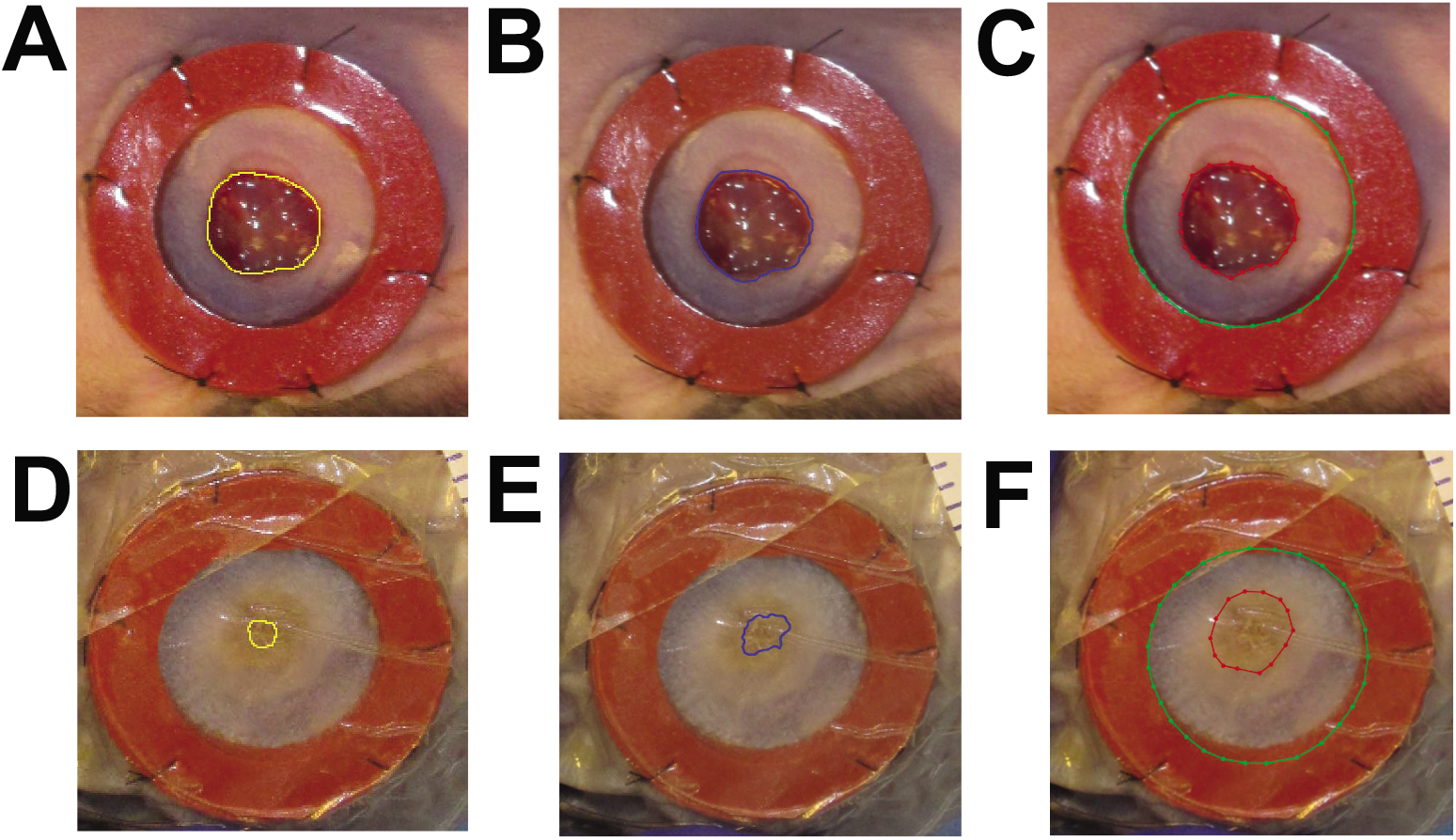
Experts vs. Periphery annotations. The left column **(A, D)** are two sample tracings performed by expert #1, middle column **(B, E)** are two sample tracings performed by expert #2, and the right columns **(C, F)** are two sample tracings performed by non-expert. **(A, B, C)** show similar trends in tracing for the same wound (taken before day 3), while **(D, E, F)** show different trends in tracing for the same wound (taken after day 3).

To compare our results to those measurements assessed by experts (see “Manual Measurement (Expert)” for more details), Fig. 16 shows manual (experts’) and automatic measurements of average wound closure percentages for C1 mice (see Fig. 16 left-plot) and C2 mice (see Fig. 16 right-plot). The orange line shows the average wound closure percentage measurements based on expert #1’s annotations, the green line shows expert #2’s annotation results, and the blue line shows the average wound closure percentages based on the automatic estimation using the developed pipeline. For C1 mice, the expert measurements seem to agree and deviate from the automated measurements, while for C2, expert #1 seems to agree with the automated measurements and deviate from expert #2’s annotation results early on. However, for all measurements, the results demonstrate that even if the mean is off, the qualitative dynamics match. In the clinical field, wound closure rates are often used to predict whether a wound will heal or not [4]. Thus having a consistent approach to compare results across experiments and time points is of utmost importance. Therefore, even if quantitative measurements are not exactly aligned, the preservation of wound healing dynamics remains informative. The ground truth can remain ambiguous but if experts can universally agree on a unique set of annotations, then the algorithms can always be retrained to segment in the same way. The segmentation algorithm can only achieve what it is taught.

**Fig 16.**
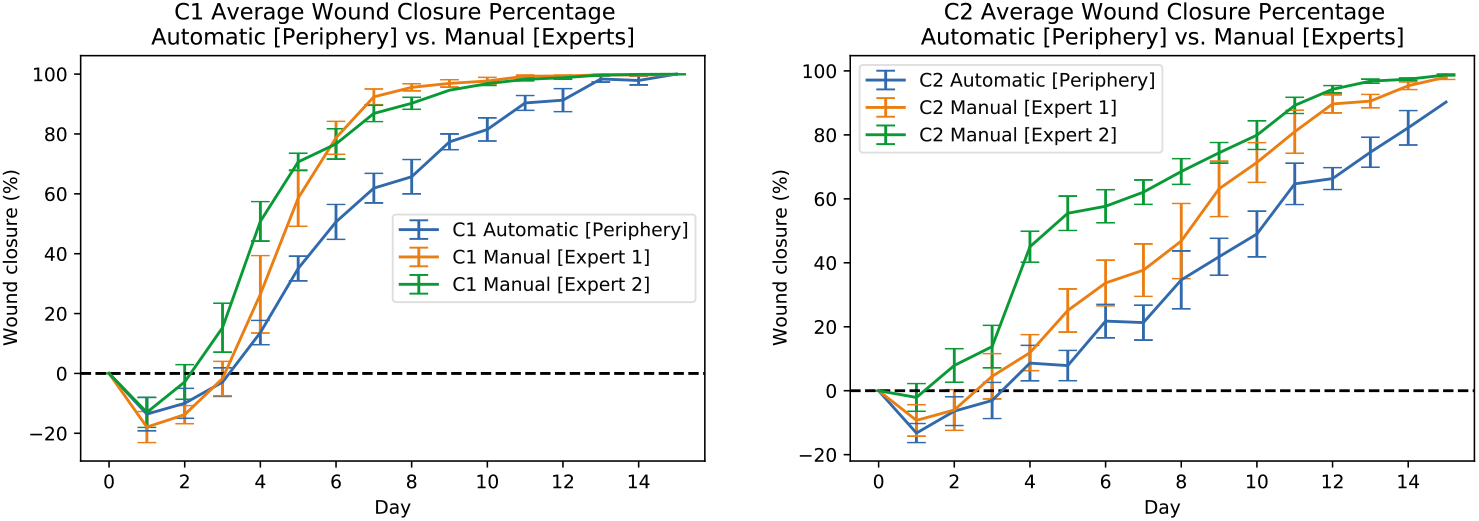
Automatic and experts measurements of wound for C1 (left-plot) and C2 (right-plot) mice. The orange line shows the average wound closure percentage measurements based on expert #1 annotations, the green line shows the expert #2 annotation results, and the blue line shows the average wound closure percentages based on the automatic estimation using the developed pipeline.

Figures (S7 Fig and S8 Fig) show the wound closure percentages for the same mice calculated from the expert #1 annotations. Figures (S9 Fig and S10 Fig) show the wound closure percentages for the same mice calculated from the expert #2 annotations.

## Conclusion

An automatic wound analysis pipeline is presented in this paper by using custom-designed deep learning-based algorithms to provide critical information such as wound closure percentages and wound size for further evaluating the wounds. The results demonstrate the effectiveness of the presented method with minimal human intervention. Considering its versatility, the developed pipeline is a promising tool for replacing the time-consuming and laborious manual process of wound measurement.

Based on the results, we observed that, on average, traces of the wound edge begin to diverge after day 3 but agreement is regained upon wound closure. There are quite a few reasons why there is some discrepancy. Human annotators take into account the previous/next images when annotating the wound edge (temporal aspect) while the model does not. Human annotators are better at dealing with some of the challenges like occlusion and blur than the model. For future work, in order to predict surface area closer to what experts measure a network could be built to incorporate contextual information such as time and specific color/texture variations. Given that defining the wound edge could be subjective and can vary from one individual to another even among experts, another interesting approach could be unsupervised methods that do not require such annotations. We postulate that the above mentioned approaches can solve the current difficulty and subjectivity by finding a universally “agreed” upon segmentation for training for a universally consistent approach to segmentation. By universally agreed on segmentation we mean a segmentation that all experts can accept.

## Supporting information

Supplemental File for Main Manuscript

## Data Availability

All data and codes necessary to replicate this study will be available here (Data+Codes) after the publication.

## Acknowledgments

Research was sponsored by the Office of Naval Research and the DARPA Biotechnologies Office (DARPA/BTO) and was accomplished under Cooperative Agreement Number DC20AC00003. The views and conclusions contained in this document are those of the authors and should not be interpreted as representing the official policies, either expressed or implied, of the Office of Naval Research and the DARPA Biotechnologies Office (DARPA/BTO) or the U.S. Government. The U.S. Government is authorized to reproduce and distribute reprints for Government purposes notwithstanding any copyright notation herein.

Funding for the BD2K Biomedical Data Science program at CSUMB is supported by the Office Of The Director (OD) of the National Institutes of Health under grant number R25MD010391, at UPR under R25MD010399, and at UCSC under NIH-1U54HG007990. The content is solely the responsibility of the authors and does not necessarily represent the official views of the National Institutes of Health.

## Supporting information

**S1 Fig. Wound closure percentages based on the automatic estimation using the developed pipeline for C1 mice both left and right wounds.**

**S2 Fig. Wound closure percentages for C1 mice both left and right wounds obtained from the manual periphery measurements.**

**S3 Fig. Wound closure percentages for C2 mice both left and right obtained using the developed pipeline.**

**S4 Fig. Wound closure percentages for C2 mice both left and right wounds obtained by using the measurements based on the manually annotated images.**

**S5 Fig. Automatic analysis of wound periphery for wounds with changed camera angle by** 10 − 15 **degrees using developed pipeline.**

**S6 Fig. Comparison of wound periphery measurements in day**0 **and day**1 **for wounds in the original dataset using developed pipeline.**

**S7 Fig. Wound closure percentages for C1 mice both left and right wounds obtained by using the measurements based on the expert** #1 **annotations.**

**S8 Fig. Wound closure percentages for C2 mice both left and right wounds obtained by using the measurements based on the expert** #1 **annotations.**

**S9 Fig. Wound closure percentages for C1 mice both left and right wounds obtained by using the measurements based on the expert** #2 **annotations.**

**S10 Fig. Wound closure percentages for C2 mice both left and right wounds obtained by using the measurements based on the expert** #2 **annotations.**

**S1 File. Measurements for all the different windows (“Error Results Ring Elimination.csv”).**

## Notes

### Competing Interest Statement

The authors have declared no competing interest.

### Summary of Updates

N/A.

