## Supplemental File for Main Manuscript for "Automatic wound detection and size estimation using deep learning algorithms"

### Supporting information

**S1 Fig. Wound closure percentages based on the automatic estimation using the developed pipeline for C1 mice both left and right wounds.** The left plot shows the wound closure percentages for the wound on the left side of 4 different C1 mice from day 0 to day 15 in different colors, and the right plot shows the wound closure percentages for the wound on the right side of those mice.

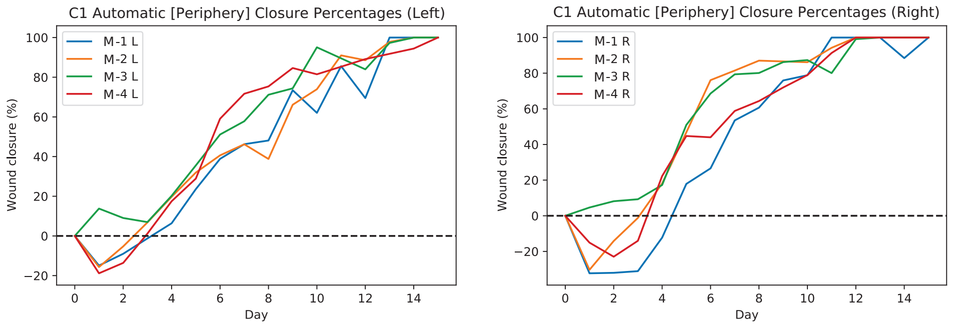


**S2 Fig. Wound closure percentages for C1 mice both left and right wounds obtained from the manual periphery measurements.** The left plot shows the wound closure percentages for the wound on the left side of 4 different C1 mice from day 0 to day 15 in different colors, and the right plot shows the wound closure percentages for the wound on the right side of those mice.

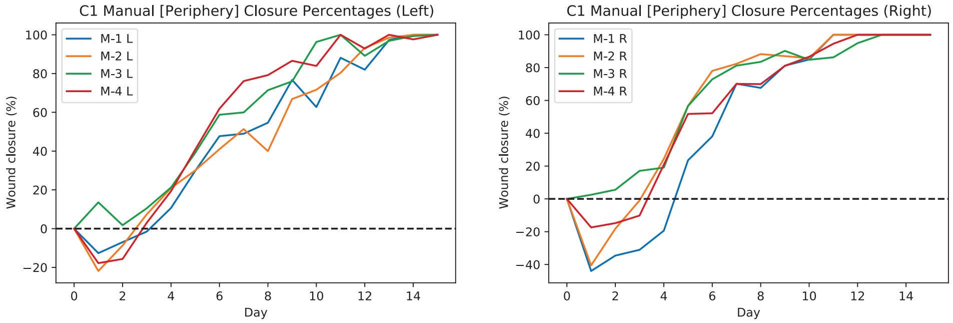


**S3 Fig. Wound closure percentages for C2 mice both left and right obtained using the developed pipeline.** The left plot shows the wound closure percentages for the wound on the left side of 4 different C2 mice from day 0 to day 15 in different colors, and the right plot shows the wound closure percentages for the wound on the right side of those mice.

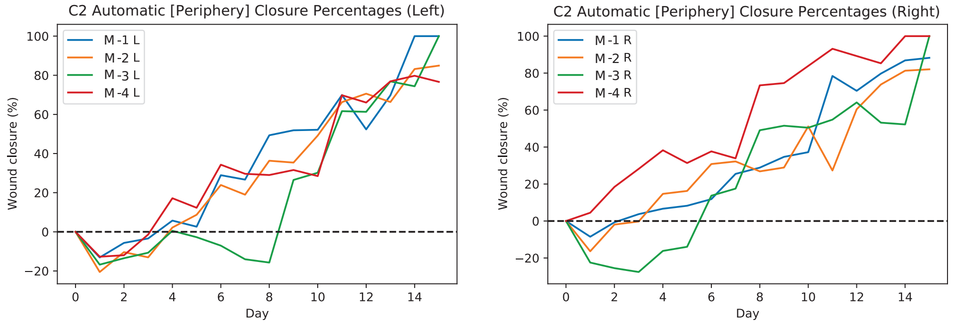


**S4 Fig. Wound closure percentages for C2 mice both left and right wounds obtained by using the measurements based on the manually annotated images.** The left plot shows the wound closure percentages for the wound on the left side of 4 different C2 mice from day 0 to day 15 in different colors, and the right plot shows the wound closure percentages for the wound on the right side of those mice.

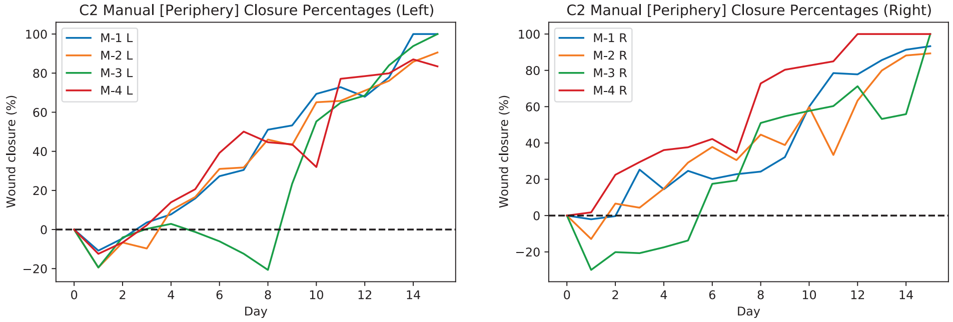


**S5 Fig. Automatic analysis of wound periphery for wounds with changed camera angle by $10-15$ degrees using developed pipeline.** The plot shows Root

Mean Square Error (RMSE) of estimated size of angled images comparing to those taken at 0 degree where the mean of the RMSE for all mice is 1.8088 and the median is 1.3472.

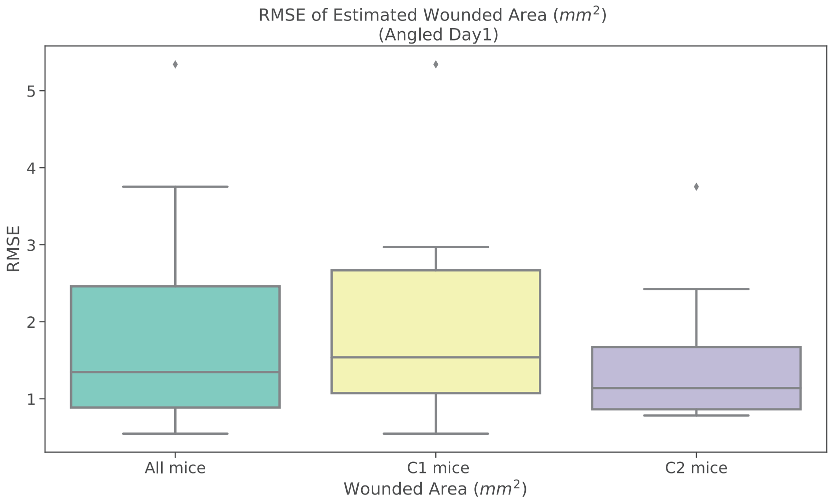


**S6 Fig. Comparison of wound periphery measurements in day0 and day1 for wounds in the original dataset using developed pipeline.** The plot shows the RMSE of images (in the original dataset) taken at Day 1 comparing to those taken at Day 0 where the mean of the RMSE for all mice is 4.1746 and the median is 4.275.

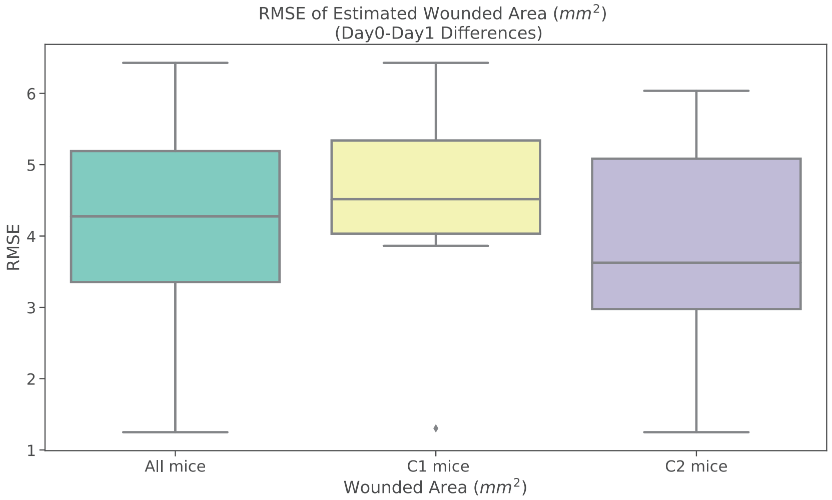


**S7 Fig. Wound closure percentages for C1 mice both left and right wounds obtained by using the measurements based on the expert #1 annotations.** The left plot shows the wound closure percentages for the wound on the left side of 4 different C1 mice from day 0 to day 15 in different colors, and the right plot shows the wound closure percentages for the wound on the right side of those mice.

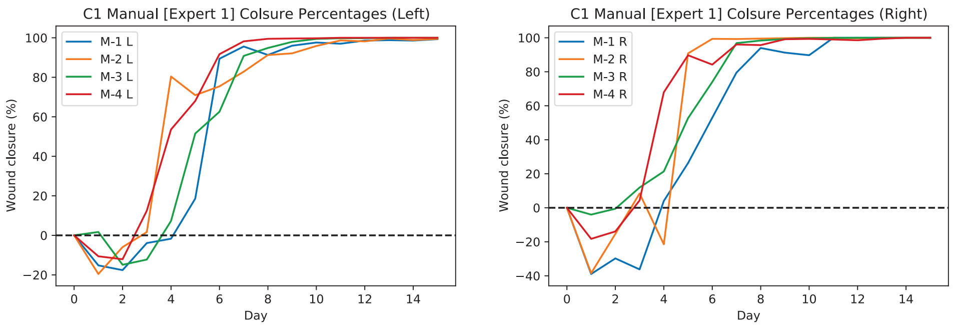


**S8 Fig. Wound closure percentages for C2 mice both left and right wounds obtained by using the measurements based on the expert #1 annotations.** The left plot shows the wound closure percentages for the wound on the left side of 4 different C2 mice from day 0 to day 15 in different colors, and the right plot shows the wound closure percentages for the wound on the right side of those mice.

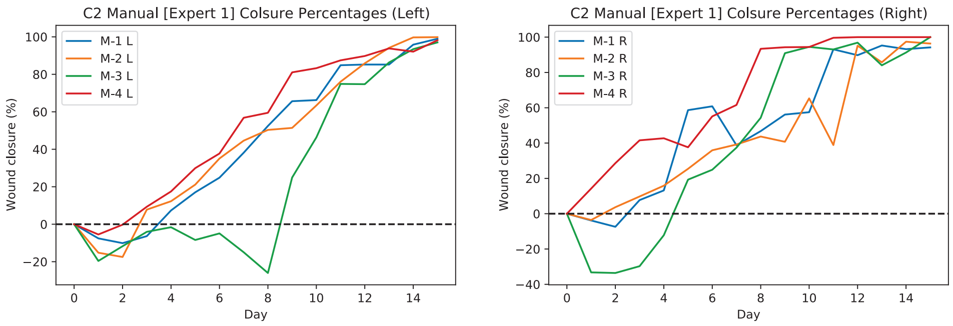


**S9 Fig. Wound closure percentages for C1 mice both left and right wounds obtained by using the measurements based on the expert #2 annotations.** The left plot shows the wound closure percentages for the wound on the left side of 4 different C1 mice from day 0 to day 15 in different colors, and the right plot shows the wound closure percentages for the wound on the right side of those mice.

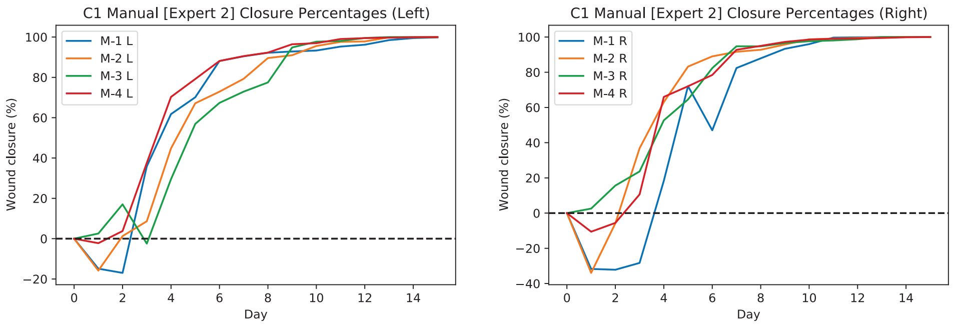


**S10 Fig. Wound closure percentages for C2 mice both left and right wounds obtained by using the measurements based on the expert #2 annotations.** The left plot shows the wound closure percentages for the wound on the left side of 4

different C2 mice from day 0 to day 15 in different colors, and the right plot shows the wound closure percentages for the wound on the right side of those mice.

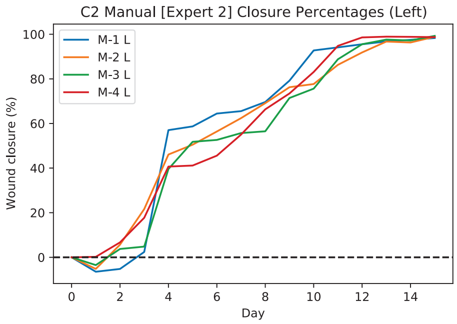


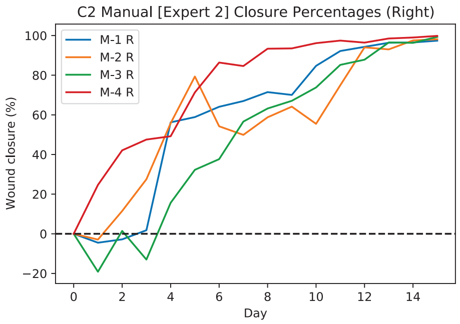


**S1 File. Measurements for all the different windows ("Error Results Ring Elimination.csv").**
